# The efficacy of sexual selection under environmental change

**DOI:** 10.1101/283457

**Authors:** Ivain Martinossi-Allibert, Claus Rueffler, Göran Arnqvist, David Berger

## Abstract

Sexual selection can promote adaptation if sexually selected traits are reliable indicators of genetic quality. Moreover, stronger sexual selection in males, as often reported in empirical studies, may help purge deleterious alleles at a low cost to population productivity. However, to what extent this remains true when a changing environment affects sexual selection dynamics has been debated. Here, we show that even if sexually selected traits remain honest signals of male quality in new environments, the efficacy of sexual selection will often be reduced under stress. We model the strength of sex-specific selection under different levels of environmental stress in a population in which males compete with each other for fertilization success and in which females experience fecundity selection. We observe that the strength of sexual selection is reduced relative to fecundity selection, resulting in a lowered potential for selection on males to aid adaptation under environmental change.

## Introduction

The question of how sexual selection contributes to local adaptation and how it aligns with the forces of natural selection can be traced back to the early days of evolutionary biology. Darwin (1871) pondered on how extreme morphological features and behaviors that seemed costly to survival had evolved, and ascribed them to selection arising from competition for mating opportunities. Later, Fisher (1958) proposed that exaggerated ornaments may become genetically correlated with female mate preferences, making the ornament and the preference coevolve in a “run-away” fashion. This view was developed further with the argument that sexual signals should be costly and only afforded by individuals of high genetic quality, leading to the “good genes” hypothesis of sexual selection (Zahavi 1975; Hamilton & Zuk 1982; Maynard-Smith 1991). According to this hypothesis, sexually selected traits are indicators of overall genetic quality because their expression reflects polygenic variation across large parts of the genome (Rowe & Houle 1996). This logic does not only apply to mate-choice signals, but also to life history, morphological and behavioral traits involved in intrasexual competition and mate searching (Whitlock & Agrawal 2009). Consequently, sexual selection favoring males of high genetic quality could increase the frequency of alleles with positive effects on female fecundity and viability (Rowe & Houle 1996; Lorch *et al*. 2003; Tomkins *et al*. 2004).

In the majority of polygamous species sexual selection acts more strongly in males (Andersson 1994; Arnqvist & Rowe 2005; Whitlock & Agrawal 2009; Janicke *et al*. 2016). As a consequence, selection in males has the potential to purge generally deleterious alleles from the population while leaving females, who experience weaker selection, relatively spared of the cost of adaptation (Manning 1984; Agrawal 2001; Siller 2001). Indeed, under the assumption that population productivity relies mainly on female fecundity, Agrawal (2001) and Siller (2001) showed that genetic load on population fitness at mutation-selection balance is inversely proportional to the degree of male-bias in selection, and therefore potentially lower in sexual relative to asexual lineages. Male-biased sexual selection for “good genes” could therefore be expected to aid adaptation in sexually reproducing species experiencing rapidly changing environments.

The effect of environmental change on sexual selection has just begun to be explored and little is known about how the benefits of sexual selection discussed above may unfold in novel environments. Empirical studies have often detected temporal and spatial variation in sexual selection metrics such as the Bateman gradient and variance in reproductive success, patterns that are frequently ascribed to ecological factors (Arnqvist 1992; Gosden & Svensson 2008; Serbezov *et al*. 2010; Byers & Dunn 2012; Robinson *et al*. 2012). Theory suggests that novel environments can perturb the expression sexual signals (for example through availability of a new ressource Kokko and Heubel 2007; Higginson and Reader 2009), although some traits actually have the potential to remain indicators of good genes across environments (for example: success in combat and mating call frequency, that are often directly related to size and therefore to condition Greenfield and Rodriguez 2004).

In this study, we develop predictions on the effect of environmental stress on sexual selection, and we find that even if the mate-choice mechanism remains unaltered, the efficiency of sexual selection for promoting adaptation is generally reduced. We employ mathematical modelling to assess the potential for sexual selection to aid adaptation by comparing the strength of selection in males and females following a rapid environmental change. In our model, males are affected by viability selection and subsequently by sexual selection resulting from competition for access to females. Females are subject to the same viability selection as males and then experience fecundity selection. Both juvenile survival and adult reproductive success is dependent on the condition of the individual. Condition, in turn, is determined by an underlying quantitative trait that was under stabilizing selection in the ancestral environment, but whose optimum has been displaced as a result of the environmental change. This causes directional selection on the trait in the novel environment. In contrast to female reproductive success, the reproductive success of a male depends on his own condition as well as on the condition of his direct competitors. Thus, sexual selection in males is frequency-dependent while fecundity selection in females is frequency-independent. To estimate the sex-bias in selection we calculate the strength of selection ***I*** (i.e., the variance in relative fitness, Crow 1958) for each sex and compare the ratio ***I***_M_/***I***_F_ for different mating systems and degrees of environmental stress. ***I*** is a theoretically appealing measure in this context because its phenotypic component provides an upper-limit to the strength of selection on any trait (independent of whether selection is directional, stabilizing, disruptive or correlational; Arnold 1986; Jones 2009), and its additive genetic component gives the per generation increase in fitness itself (Fisher 1930; Houle 1992). In our model, the only source of variation in fitness is the underlying quantitative trait (there is no environmental variance), therefore ***I*** directly represents the strength of selection. ***I*** has also been used extensively in studies of both natural and laboratory populations (Janicke et al. 2016). Thus, using ***I*** makes our model accessible to empiricists while providing testable predictions. We provide an analysis of empirical data in two species of seed beetle to illustrate its utility. We also compare ***I***_F_, the response of female fitness to selection in females, to COV_MF_, the intersexual genetic covariance for fitness, which represents the response in female fitness to selection on males. This comparison allows us to determine whether sexual selection on males can contribute to the recovery of female fitness following a rapid environmental change.

We find that, as expected, both ***I***_M_ and ***I***_F_ increase as the environment changes and becomes stressful. However, the increase in ***I***_M_ under environmental change is critically dependent upon mating system parameters, such as the number of individuals competing (lek size) and the degree to which differences in condition between competing males affects fertilization success (g). We refer to *g* as the skew parameter (cf. Kokko and Lindström 1997). For lek sizes relevant for most natural systems, the potential benefits of sexual selection (***I***_M_ ***/ I***_F_) decreases with increasing environmental stress. Comparing the ratio COV_MF_ / ***I***_F_ gives qualitatively similar results, confirming that the relative efficacy of good genes sexual selection is reduced in populations subject to a rapid environmental change.

## Model

### The population

The model developed in this study quantifies the strength of sex-specific selection in a sexually reproducing population that is exposed to stress by a sudden change in environmental conditions. We consider an organism with non-overlapping generations.

Individuals are characterized by a quantitative trait *z* with frequency distribution *f*_*z*_ in a population. Both sexes have the same trait distribution, i.e., there is no sexual dimorphism. As a result of a long period of stabilizing selection, trait values in the population are normally distributed with mean *μ*_*z*_ *= z*_*0*_ and standard deviation *σ*_*z*_. For a trait value *z* we thus obtain

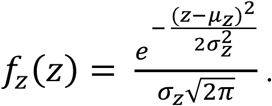

The match between the trait value *z* and the environmentally determined optimal trait value *z*_*0*_ determines an individual’s condition *c*. We assume that *c* is bound between 0 and 1 and that the decrease in condition with increasing deviation from *z*_*0*_ follows a Gaussian curve. Thus

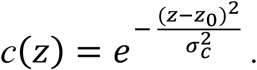

Here, *σ*_*c*_ determines the width of the Gaussian curve and smaller values imply that the condition decreases faster with increasing distance between *z* and *z*_*0*_. The phenotypic standard deviation of quantitative traits is typically on the order of 1/5 to 1/100 of this width-parameter when estimated from studies of stabilizing selection (Johnson & Barton 2005). We model all scenarios using these two extreme values (*σ*_*z*_ /*σ*_*c*_ = 1/5 and *σ*_*z*_ /*σ*_*c*_ = 1/100). However, changing the amount of standing variation in *z* has little effect on the results and we therefore present only those for *σ*_*z*_ /*σ*_*c*_ = 1/5. From the distribution of trait values *f*_*z*_ and the function *c(z)*, we obtain for the distribution of conditions *f*_*c*_ within a population

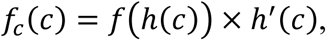

where *h*(*c*) is the inverse function of *c*(*z*) and *h*′(*c*) its derivative (e.g. Weisstein 2003).

### Survival to reproduction

The probability *s* to survive to sexual maturity is determined by the condition *c* according to the monotonically increasing function

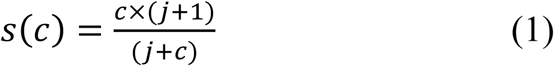

(Figure 2). Here, the parameter *j* determines the intensity of viability selection. For *j=0* there is no viability selection and all individuals survive to reproduction. For *j=0* we obtain *s*(*c*)=1 for all values of *c* and for *j* positive we obtain *s*(0)=0 and *s*(1)=1, and survival increases from 0 to 1 with increasing condition. As *j* increases the relationship between *c* and *s* changes from a saturating function toward a linear function (Figure 2b). We present results for the three levels of viability selection shown in Figure 2b: *j=*0 (no viability selection), *j=*0.1 (weak viability selection) and *j=*10 (strong viability selection). The distribution of *c* in the population after viability selection and before reproduction in both sexes is given by

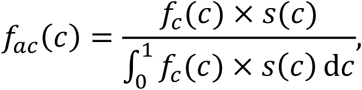

where the numerator is a standardization factor such that *f*_*ac*_ is indeed a probability density function.

**Figure 1.**
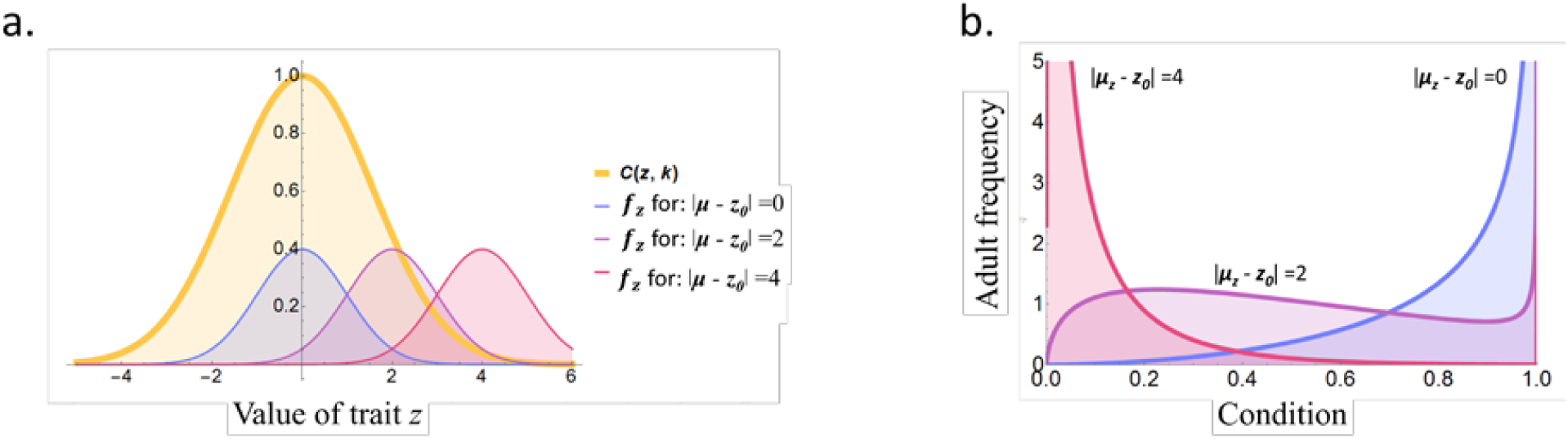
Effect of environmental change on the probability density function (PDF) of (a) trait *z*, and (b) condition *c*. In (a) the yellow Gaussian curve represents the relationship between trait value and condition, and PDFs for *z* are given for three levels of environmental change: |*μ*_*z*_ *- z*_*0*_|=0 (blue curve), |*μ*_*z*_ *- z*_*0*_|=2 (purple curve) and |*μ*_*z*_ *- z*_*0*_|=4 (pink curve). In (b) the PDFs of condition *c* resulting from mapping the trait to condition (yellow curve in (a)) are given for the three levels of environmental change.

**Figure 2.**
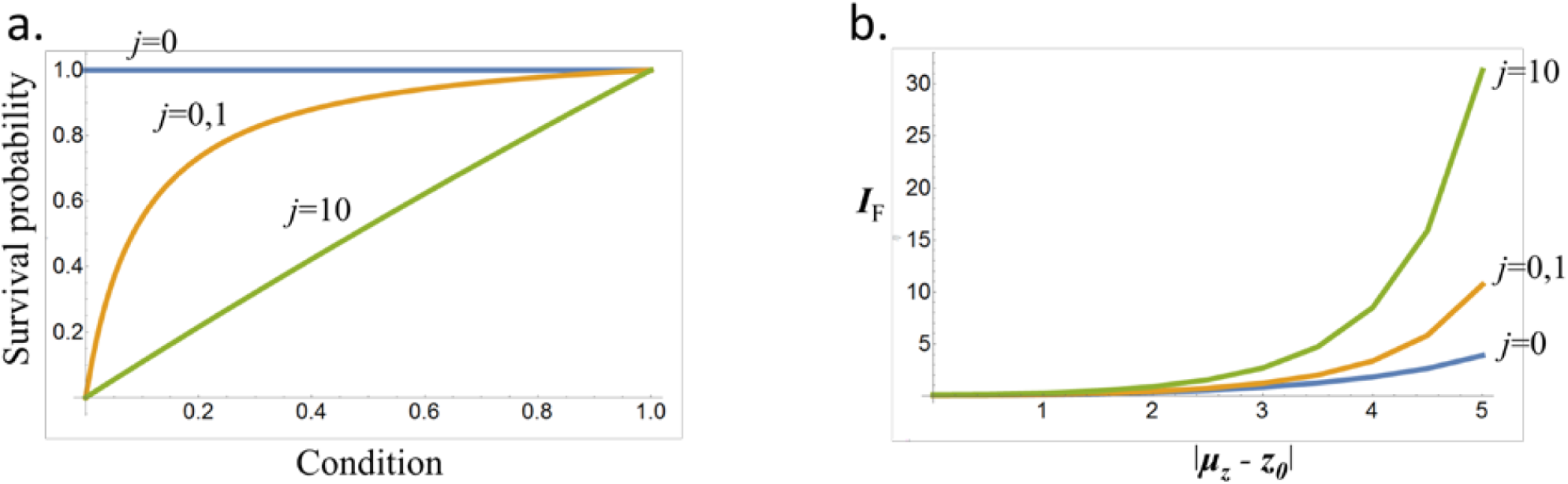
Effect of parameter *j* on (a) the relationship between condition and survival probability *s* and (b) the opportunity for selection in females as the population is exposed to environmental change. Environmental change is given by the distance between optimal trait value and population mean (i.e., |μ - z_0_|) in units of standard deviation of the Gaussian fitness landscape.

### Male selection

Reproductive success is the outcome of competition for access to females and determined in a group of *n* competing males. For simplicity, we refer to this group as a lek throughout, although our model can represent many different types of mating systems as will be discussed below. The proportion of females fertilized by a focal male with condition *c*_*1*_ in a lek with *n-1* competitors with conditions ***c*** = (*c*_*2*_,*…,c*_*n*_) is given by

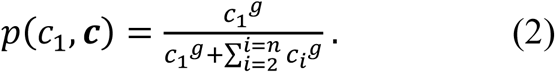

Here, the positive constant *g* determines the degree to which differences in condition among competing males results in differences in fertilization success. With *g* = 0 fertilization success is independent of male condition and there is no sexual selection on condition. When *g* = 1, males have a fertilization success equal to the proportion of their condition to total condition within the lek. This scenario could, for example, represent a sessile broadcast spawning organism for which reproductive success is proportional to the investment in gamete production. With increasing *g* fertilization success becomes increasingly skewed towards the individual with the highest condition in the lek and in the limit of *g* = ∞ the single best individual fertilizes all offspring. High values of *g* can be thought of as a mating system where males compete for full control of fertilization in a harem of females, such as often suggested for elephant seals or lions. We refer to *g* as the skew parameter (cf. Kokko and Lindström 1997)

Male fertilization success *m* is the proportion *p* of fertilized females multiplied with the number of available females. Assuming an even sex-ratio in a lek this number equals *n*. Thus,

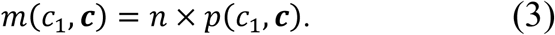

On average, fertilization success equals 1 (on average, one male fertilizes one female as a result of an even sex-ratio; Fisher 1930, Houston and McNamara 2005, Kokko and Jennions 2008), and the highest possible reproductive success of a male in a lek of size *n* equals *m=n*. There is no assortative mating and males mate with a randomly drawn set of females.

For each value of *z* the expected value of *m* in a given population can be calculated by integrating over the possible conditions of all competitiors in a lek. However, solving this integral analytically proved unfeasible for larger leks and a numerical approximation described in the Appendix 1 is used.

Combining equations (1) and (3) allows us to calculate male fitness *w*_m_, which is the product of survival probability *s* and reproductive success *m*. Thus,

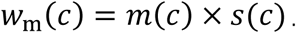

The mean and variance in male fitness are then approximated with another numerical procedure described in the Appendix 1. This allows us to calculate the opportunity for selection on males (variance in relative fitness),

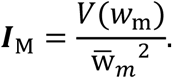

### Female selection

Female fecundity is assumed to be linearly increasing with condition *c*. Female fitness *w*_f_ is then the product of the probability to survive to sexual maturity, given by equation (1), and fecundity. Thus

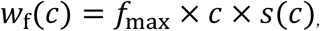

where *f*_max_ is the maximal possible fecundity. The mean and variance in female fitness is determined by a similar procedure as described for male fitness (see Appendix 1). The opportunity for selection in females (variance in relative fitness) is

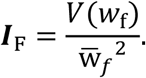

### Male-Female covariance in fitness

Similarly to ***I***_M_ and ***I***_F_, COV_MF_ is calculated using a numerical procedure described in the Appendix 1.

### Environmental stress

In the ancestral environment the population mean trait value *μ*_*z*_ equals the trait value that maximizes condition, *μ*_*z*_*=z*_*0*_. Environmental change is modelled by moving the optimal trait value *z*_*0*_ away from *μ*_*z*_ while keeping the strength of stabilizing selection (determined by the parameter *σ*_*c*_) constant. With increasing difference |*μ*_*z*_ *- z*_*0*_| the condition *c* decreases for the majority of the population. The distribution of conditions in the population is affected as follows (see Figure 1). In the ancestral environment (|*μ*_*z*_ *- z*_*0*_| = 0) the distribution of conditions is skewed towards high values close to 1 as most individuals in the population are well-adapted. At intermediate stress (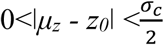, resulting in a reduction in mean reproductive success of up to 35%) the density distribution of conditions is more symmetric and uniform. Finally, under high stress (|*μ*_*z*_ *- z*_*0*_| >*σ*_*c*_/2, resulting in a decrease in reproductive success between 40 to 90%) the distribution of conditions becomes skewed towards low values close to 0.

## Results

### Selection in the ancestral environment

The strength of selection in males ***I***_M_ (and the resulting ratio ***I***_M_ / ***I***_F_) increases with the skew parameter g. Second, a larger lek size *n* provides a chance for the best males to monopolize a larger number of females, which increases ***I***_M_. However, ***I***_M_ is a saturating function of *n* and the reproductive success of the best male rarely reaches the absolute maximum set by lek size, as can be seen from Figure 3b. For low values of *g* (i.e., differences in male conditions affect fertilization success only weakly) this saturation starts already at a rather low lek size because males are not able to monopolize all the females in the lek. At higher values of *g* saturation is reached only for larger leks as high condition males can monopolize a larger number of females in this type of mating system (Figure 3a). Intuitively, increased viability selection makes the strength of selection more similar in the sexes.

**Figure 3.**
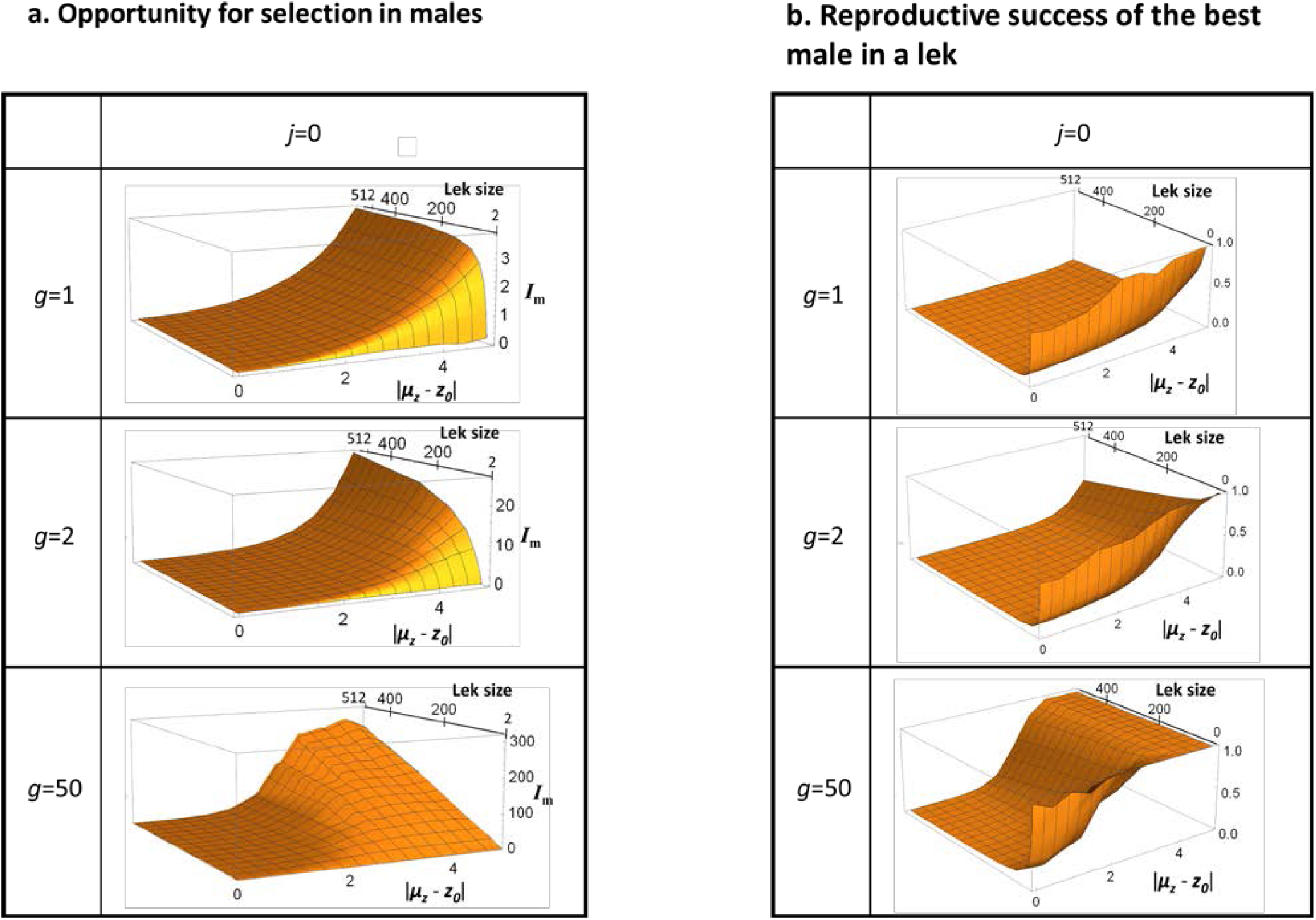
(a) Male opportunity for selection and (b) reproductive success of the best male as a function of environmental change and lek size, presented for several values of the skew parameter *g*. The reproductive success of the best male in (b) is presented as the proportion of fertilized females relative to the total number of females in the lek to make it more comparable across different lek sizes. The absolute limit to male reproductive success is equal to lek size, and the ratio presented in (b) equals 1 when this limit is reached.

### Selection following environmental change

Environmental stress increases directional selection on *z*. This results in increased strength of selection ***I*** in both sexes (see Figure 2b and 3a). However, the effect of stress also interacts with characteristics of the mating system. In figure 4 we summarize the effect of environmental stress on the relative strength of selection in males and females (log_*e*_(***I***_M_ ***/ I***_F_) for different mating systems (combinations of parameters *<u>g</u>* and *n*) for the three different intensities of viability selection. As a baseline scenario we consider *g=1*. In this case the fertilization success of a focal male equals the proportion of its condition relative to the sum of conditions of all males in the lek. This scenario serves as a reference point because adult selection then operates in a similar way in males and females; reproductive success in females equals the ratio of female fecundity over the population’s average. The only difference between the sexes is that selection in females acts across the whole population, while males compete to fertilize only the eggs that are available locally within a lek. Consequently, with *g=*1, males and females experience the same strength of selection when lek size becomes infinite (i.e., the lek becomes the entire population). This effect can be observed for large lek sizes (Figure 4, *g*=1). For a lek size of 512 the log ratio ***I***_M_ ***/ I***_F_ is very close to 0 (i.e. the strength of selection is equal in the sexes).

**Figure 4.**
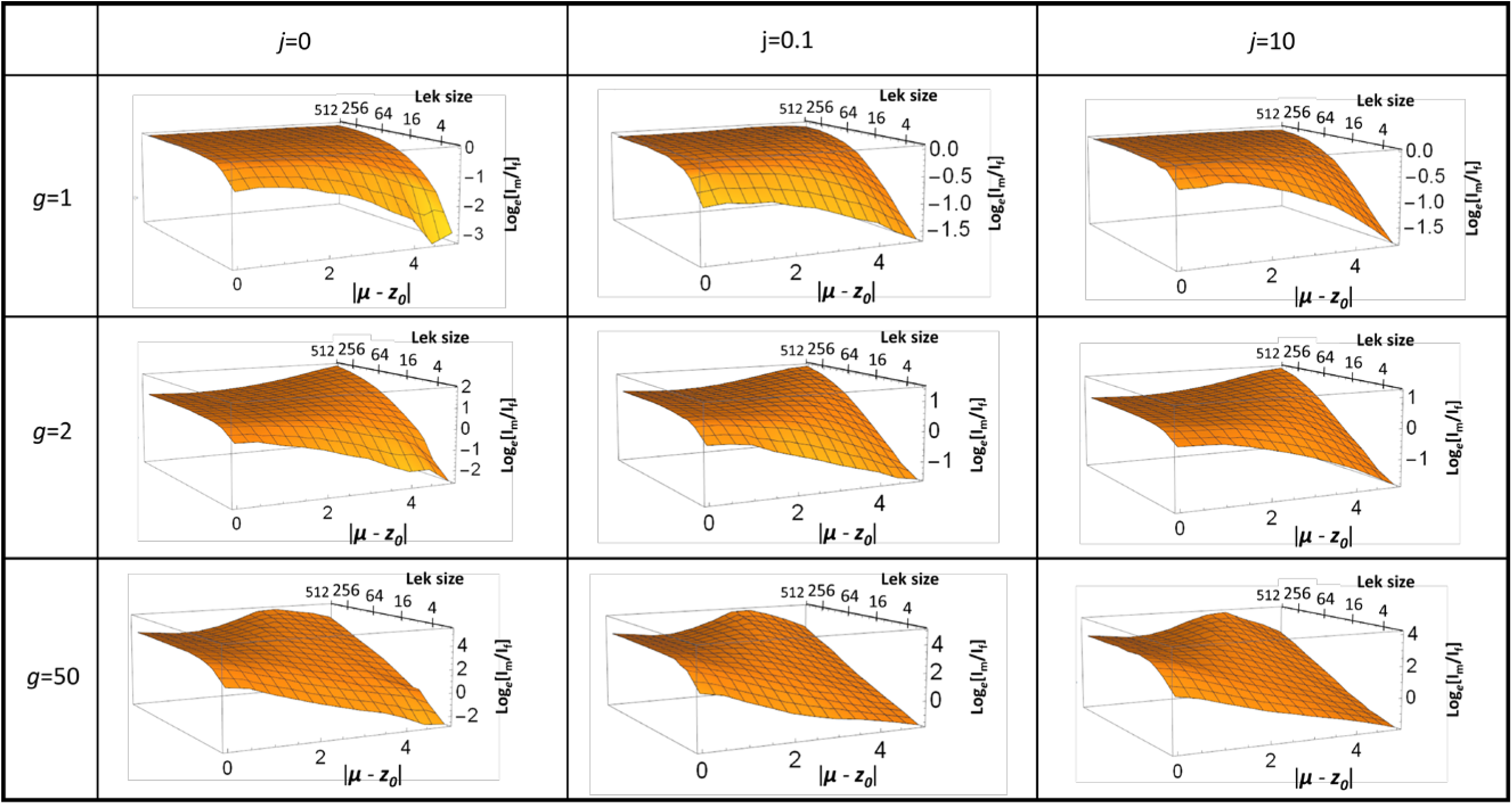
Log_e_-ratio of male over female strength of selection as a function of lek size and environmental change for three different values of the skew parameter *g* (rows) and three different strengths of survival parameter *j* (columns). Environmental change is represented by the distance between optimal trait value and population mean (|μ - z_0_|) in units of standard deviation of the Gaussian fitness landscape. Lek size is plotted on a log_2_-scale.

For small leks, increasing stress decreases the ratio ***I***_M_ ***/ I***_F_ independently of the other model parameters. This is because male reproductive success is bounded by the absolute maximum number of females that the best male can monopolize (which is equal to the size of the lek), and this sets an upper boundary to the variance in male reproductive success. However, for larger leks stress either uniformly increases the ratio ***I***_M_ ***/ I***_F_ weakly (observed for *g=2*), or first increases this ratio and then decreases it at very high levels of stress (observed for *g=50*). To understand these patterns it is helpful to inspect the reproductive success of the best male, presented in Figure 3b. First, males in well-adapted populations never monopolize a large proportion of the females in the lek, regardless of the value of the skew parameter *g*. The reason is that in benign environments most individuals are in good condition, and therefore even the highest condition males will frequently encounter almost equally good competitors. This phenomenon explains why ***I*** is never extremely male-biased in well-adapted populations (Figure 4). Second, the maximum reproductive success is for most parameter combinations lower than lek size, even under high levels of stress, which means that even the best male in a lek very rarely can father all offspring. For *g*>1, stress tends to increase the maximum male reproductive success toward lek size (Figure 3). This is because high condition individuals become rarer as stress increases, allowing such individuals to monopolize females in their lek because their competitors are of comparatively low condition. This phenomenon is magnified by increasing *g* (Figure 3b). This mechanism explains why selection becomes more male-biased under stress for high lek sizes as can be seen in Figure 4 for *g*=2.

Finally, the observation that for *g*=50 the male-bias in selection is maximal at intermediate stress and then decreases at higher stress can be understood based on Figure 3b. Due to the extreme intensity of competition, high condition males can in this case monopolize all females in the lek already before extreme levels of stress are reached and therefore the most extreme value of reproductive success ceases to increase after intermediate stress is reached. As variance in female reproductive success is directly proportional to variance in condition *c* (which increases with environmental stress), this results in a decreasing ratio of ***I***_M_ ***/ I***_F_ from intermediate to high stress.

The ratio COV_MF_/***I***_F_ responds to environmental change and to the mating system characteristics in qualitatively similar ways as the ratio ***I***_M_ ***/ I***_F_ (Appendix 2), thus confirming that the efficacy of good genes sexual selection is generally reduced relative to fecundity selection following environmental change.

## Discussion

Environmental change will render many natural populations maladapted to their novel ecological settings. Here, we investigated by means of a mathematical model to what extent “good genes” sexual selection can facilitate rapid adaptation under such scenarios. Our approach is motivated by previous theoretical findings showing that sexual reproduction can offer a population-level benefit if purifying selection is stronger in males than in females, sparing the latter of the cost of adaptation (Manning 1984, Agrawal 2001, Siller 2001, Whitlock & Agrawal 2009). Thus, to evaluate the role of sexual selection in aiding adaptation, we compared the strength of selection in males competing over fertilization success to that of fecundity selection in females across different levels of population maladaptation to local ecological conditions.

We find that while environmental change increases selection in both males (***I***_M_) and females (***I***_F_), the effect on the ratio ***I***_M_/***I***_F_ depends on the size of the lek (i.e., the number of males that compete for matings over the same number of females) and the skew parameter *g*, which determines the degree to which differences in condition among competing males results in differences in fertilization success. For small lek sizes the ratio ***I***_M_/***I***_F_ decreases strongly with environmental stress, indicating that the efficiency of sexual selection is reduced under stress. This effect occurs because lek size imposes a limit on variance in male reproductive success, which thus limits the increase of variance in male reproductive success with stress. For large leks, ***I***_M_/***I***_F_ remains qualitatively unchanged or even can increase with environmental stress. The latter effect happens because high condition males become rare under high stress and these males can then monopolize a large number of females within their lek, resulting in a strong increase in ***I***_M_. These findings highlight the interplay between mating system characteristics and ecological conditions in deciding evolutionary demography in sexually reproducing species.

Our results show that the benefit of sexual selection through competition among males can be reduced by a rapid environmental change if males compete in groups below a certain size (here approximately 30, although we expect this number to depend on finer ecological details). We suggest that this scenario best represents sexual selection in nature because in most natural populations the maximum number of females that a male can monopolize is likely not very high. Indeed, even if a male can achieve a status of dominance over other males, there should be a limit to the number of females this male can guard in order to secure paternity of all their offspring. We can make an attempt at estimating the maximum number of partners that a male can monopolize across its lifetime using data from natural populations; for example, in the long-term data of Soay sheep on the island of St. Kilda the most successful male ever recorded since the 1960s, OG023, sired around 40 lambs (http://soaysheep.biology.ed.ac.uk/). If we consider that adult females reproduce on average during five breeding seasons and produce one offspring per season, OG023 monopolized the reproductive success of 8 females throughout his lifetime, making the upper limit for the “lek size” (corresponding to our model parameter *n*) of this population 8. Another example are field crickets studied by Rodriguez-Muñoz et al. (2010), where a similar calculation indicates that the most successful male monopolized a bit more than 9 females over its lifetime. Data from a long-term study in the collared flycatcher (Merilä & Sheldon 2000) suggests that the best male recorded monopolized some 9 females. Finally, field data on the elephant seal, a harem species, suggests that alpha males may be able to monopolize up to 80 females during a particular breeding season (LeBoeuf 1974), but this is most likely a large overestimation of the lifetime mating success of the best male, because it is probable that the investment necessary to reach dominance in such systems can be achieved only during one breeding season over the life of a male (Clutton-Brock & Sheldon 2010). Thus, while information on natural variation in male fertilization success is scant for most species and the calculations above are rough approximations, the currently available data seem to indicate that “lek size” rarely becomes very large.

Our model assumes that female fitness relies mainly on fecundity and male fitness being mainly the outcome of competition for matings. These are simplifying assumption because in most species females also compete for resources and mating opportunities to some extent, and males vary in fertility. Take, for example, the studies we used above to assess natural variation in “lek size”. Male and female flycatchers have similar variance in reproductive success and maximum reproductive success (Merilä and Sheldon 2000) and in the study of field crickets (Rodriguez-Muñoz et al. 2010) both males and females benefit equally from remating (i.e., they have similar Bateman gradients) which means that sexual selection seems to acts equally in the sexes. In some species or populations, females may also compete more strongly than males do (e.g. Oring et al. 1983; Reynolds et al. 1986; Berglund and Rosenqvist 2003; Fritzsche and Arnqvist 2013). A key finding of our exploration is that frequency-dependent selection is affected differently than frequency-independent selection by environmental change. This result should hold qualitatively even if the sexes are not subject to only one type of selection, but the predictions of the model must be adapted to the ecology of the species studied. Nevertheless, sexual selection itself is an inherently frequency-dependent process, and this form of selection acts more strongly in males in most species (e.g. Janicke et al. 2016), which was the main motivation for our model.

Our model presents a snapshot of the strength of selection experienced by a population immediately after a rapid environmental change. It does not include an evolutionary response to the novel environmental conditions. However, quantitative genetic theory often make similar assumptions (e.g. constant genetic variances and trait distributions in the face of continuous directional selection) that are unlikely to be valid for longer evolutionary time frames, but may nevertheless give reasonable predictions of short term evolution from standing genetic variation (Lynch & Walsh 1998). Assuming that population demography is mainly regulated through the reproductive output of females, we can thus ask how population fitness should evolve through genetic responses in female viability and fecundity. These responses are driven by direct selection on female fitness, as well as by a genetically correlated response to selection in males. Those two elements can be calculated as the genetic variance in female fitness (***I***_F_), and the genetic covariance between male and female fitness (COV_MF_), respectively, where COV_MF_ gives the response of female fitness to selection in males. Comparing COV_MF_ to ***I***_F_ quantifies whether sexual selection on males benefits female fitness more or less than fecundity selection on females. In the present model, the responses of ***I***_M_/***I***_F_ and COV_MF_/***I***_F_ to environmental change and mating system characteristics are qualitatively identically (Appendix 2). This is expected because male and female fitness are positively correlated due to the fact that they share the same genetic architecture for trait *z*. As a result, selection in males will automatically benefit females and ***I***_M_ is a good proxy for the benefits of sexual selection. We note, however, that while the assumption that phenotypic condition shares a common genetic basis in males and females should hold true for generally deleterious mutations, it might not always be the case. For example, in situations where sexually antagonistic genetic variation is dominant, the effects of stress may differ from those modelled here (e.g. Long et al. 2012; Berger et al. 2014, Connallon and Clark 2014).

### Conclusion

The relationship between natural and sexual selection has puzzled biologists since the days of Charles Darwin. More recently, the potential good genes benefits of sexual selection have been highlighted by the genic capture hypothesis (Rowe and Houle 1996, Lorch et al. 2003, Tomkins et al. 2004) but how and if such benefits materialize under novel ecological conditions is not well understood and poses an outstanding problem in today’s times of rapid environmental change. While recent theory (Greenfield and Rodriguez 2004, Kokko and Heubel 2007, Higginson and Reader 2009, Holman and Kokko 2014) suggests that novel environments may disrupt the mate choice process by decreasing the reliability of sexual signals, putting limits to the efficacy of sexual selection and its potential to promote local adaptation, these conclusions are sensitive to precise ecological settings and thus remain highly contentious. Our model suggests that, even without changes to the mate choice process, the efficiency of sexual selection is generally reduced under environmental stress due to its frequency dependent nature. The range of natural variation in lek size is difficult to estimate (it requires life time data on genetic parentage) but we suggest that relatively small “realized” lek sizes (n<60) are the rule in animals, indicating that our predictions may apply to most organisms. Our findings further suggest that contemporary environmental change will to a variable extent hamper the importance of sexual selection for adaptation, depending on the exact characteristics of the mating system and the ecology and life history behind sex-specific reproductive competition. Nevertheless, our model provides general and testable qualitative predictions of the effect of environmental stress on adaptation through sexual selection. We have illustrated this by the analysis of our own dataset on two species of seed beetle (Appendix 3), and we hope that this will encourage further research efforts along these lines.

