## Supplementary Materials for "The efficacy of sexual selection under environmental change"

### Appendix 1

***Male variance in fitness***

Numerical simulations are used to compute male fertilization success *m* for 500 regular spaced values of condition *c* of the focal male, ranging between 0 and 1. For each of these values, a vector ***c*** containing the conditions of the *n-1* competitors was determined by randomly drawing *n-1* times from the probability density function of adult conditions. To reduce the stochasticity introduced by drawing competitors from a distribution, male fertilization success was calculated for each individual as the average over 10 randomly assembled leks.

To calculate the mean and the variance in male fitness we discretize the distribution of conditions in the adult population into 500 bins with the above determined focal *c*-values as midpoints (with 500 regular spaced *c*-values between 0 and 1 we obtain bins of width 0.002). The density of individuals with e.g. *c*=0.1 is calculated by integrating the distribution $f_c$ over the interval (0.099, 0.101). The mean and variance in male fitness are then given by

$$\bar{\mathrm{w}}_m = \frac{1}{500} \sum_{i=1}^{i=500} w_{\mathrm{m}}(c_i) \times \int_{c_i - 0.001}^{c_i + 0.001} f_c(c)\, \mathrm{d}c$$

and

$$V(w_{\mathrm{m}}) = \frac{1}{500} \sum_{i=1}^{i=500} [w_{\mathrm{m}}(c_i) - \bar{\mathrm{w}}_m]^2 \times \int_{c_i - 0.001}^{c_i + 0.001} f_c(c)\, \mathrm{d}c,$$

respectively.

***Variance in female fitness***

The mean and variance in female fitness is determined by the same procedure as described for male fitness with the exception that no replicates are necessary to calculate expected female fitness. Thus,

$$\bar{w}_f = \frac{1}{500}\sum_{i=1}^{i=500} w_\mathrm{f}(c_i) \times \int_{c_i-0.001}^{c_i+0.001} f_c(c)\,\mathrm{d}c$$

and

$$V(w_\mathrm{f}) = \frac{1}{500}\sum_{i=1}^{i=500} [w_\mathrm{f}(c_i) - \bar{w}_f]^2 \times \int_{c_i-0.001}^{c_i+0.001} f_c(c)\,\mathrm{d}c.$$

We note that the distribution of female fitness can in fact be determined without discretizing the distribution of conditions but we preferred to use the same numerical simulation technique to ease subsequent comparisons between the sexes.

***Male-Female genetic covariance for fitness***

Similarly to male and female variance in fitness, $COV_{MF}$, the male-female covariance for fitness, was calculated using numerical simulations using 500 values of condition *c*

$$COV(w_\mathrm{m}, w_\mathrm{f}) = \frac{1}{500}\sum_{i=1}^{i=500} [w_\mathrm{m}(c_i) - \bar{w}_m] \times [w_\mathrm{f}(c_i) - \bar{w}_\mathrm{f}] \times \int_{c_i-0.001}^{c_i+0.001} f_c(c)\,\mathrm{d}c.$$

| | *j*=0 | j=0.1 | *j*=10 |
|---|---|---|---|
| *g*=1 | 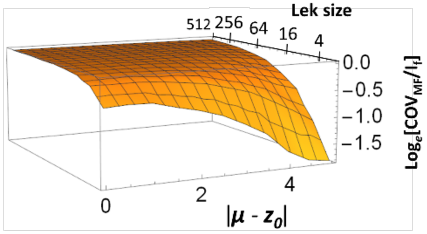 | 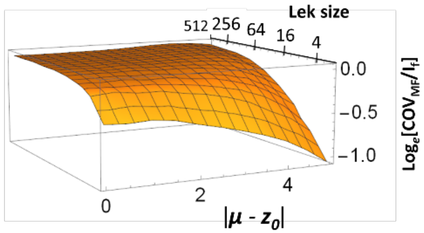 | 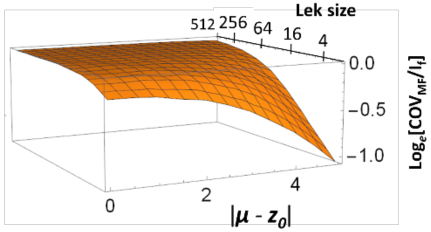 |
| *g*=2 | 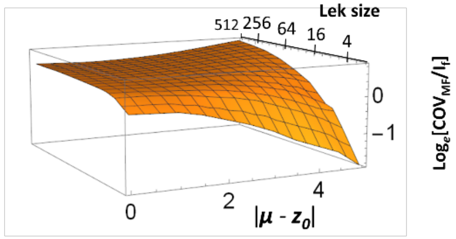 | 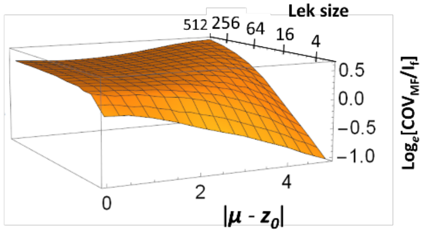 | 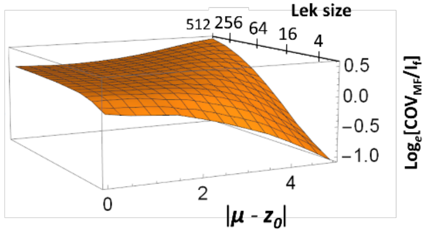 |
| *g*=50 | 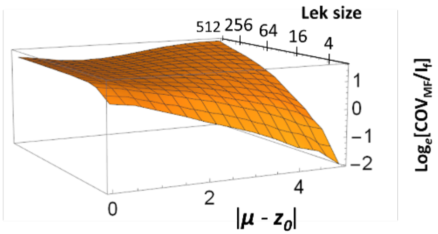 | 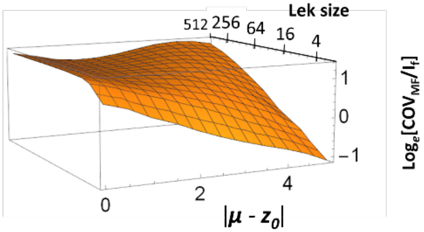 | 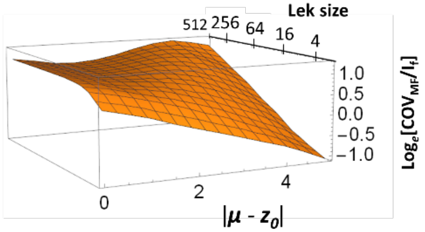 |

**Appendix 2. Log$_e$-ratio of male-female covariance over female strength of selection** presented as as a function of lek size and environmental change for three different values of the skew parameter *g* (rows) and three different values of the survival parameter *j* (columns). Environmental change is represented by the distance between optimal trait value and population mean ($|\mu - z_0|$) in units of standard deviation of the Gaussian fitness landscape. Lek size is plotted on a $\log_2$-scale.

**Appendix 3. A test of predictions at low lek-size using data on seed beetles**

To test model predictions for the smallest lek-sizes we reanalyzed data from our own experiments in the seed beetle *Callosobruchus maculatus* (Berger *et al.* 2014) and the bean beetle *Acanthoscelides obtectus* (Martinossi-Allibert et al. 2018). The experiments are described in detail in the original publications, so here we only give a brief description. In these studies, focal males and females were allowed to compete against sterilized competitors for mating with opposite sex individuals. The competitor and mating partners stem from a standard reference population of the same evolutionary background and raised in the same environment as the focal individual. All individuals were virgin and 0-24h old at the start of the assays. The individuals were allowed to mate and lay eggs until their death. Hence, the assays measured lifetime reproductive success and included both pre- and postcopulatory selection in males.

For *C. maculatus* two male competitors and three reference females were provided in the male assays, whereas focal females were provided with two reference males in the female assays. For *A. obtectus* assays were the same for males and females; focal individuals were provided with one sterilized reference competitor and two fertile opposite sex mating partners. Hence, lek-size was 2-3 in these assays. The study on *C. maculatus* estimated lifetime reproductive success across a benign (29°C) and stressful (36°C) temperature in two beetle populations isolated from central Africa (see Berger et al. 2014). In the study on *A. obtectues* the effect of host plant stress was investigated in two sets of experimental evolution lines that had been evolving for 80 generations on one or the other of two alternative hosts (see Martinossi-Allibert et al. 2018). These two sets of lines were measured for lifetime reproductive success on both hosts in a reciprocal 2x2 design. Both studies provided nearly 1000 independent estimates of reproductive success for each sex, environment and population.

Together, these two studies generated four paired comparisons of the opportunity for selection in males and females across a benign and stressful environment. To test our model predictions we ran Markov Chain Monte Carlo (MCMC) resampling using the MCMCglmm package (Hadfield 2010) in R (R core team 2013). We estimated the opportunity for selection (***I***) as the variance in relative fitness within each sex, environment and population, while controlling for block effects. Simulations were run using standard weak and uninformative priors using the idh structure. We ran 110.000 simulations for each of the four models, where the first 10.000 simulations were discarded, and saved every $100^{th}$ simulation. This yielded 1000 uncorrelated (autocorrelations <0.05) posterior estimates of ***I*** in each sex. Based on this we calculated 95% credible intervals for $\boldsymbol{I}_M$ and $\boldsymbol{I}_F$ and the ratio $\boldsymbol{I}_M/\boldsymbol{I}_F$. We then tested if the ratio $\boldsymbol{I}_M/\boldsymbol{I}_F$ was greater in benign relative to stressful environments.

Indeed, environmental stress seems to increase selection in both males and females (Fig A3.A) and decrease the ratio $\boldsymbol{I}_M/\boldsymbol{I}_F$ in all four cases (Fig. A3.B). A paired *t*-test suggests that the observed decrease of the ration $\boldsymbol{I}_M/\boldsymbol{I}_F$ is not due to chance (t = 3.78, df=3, two-sided P = 0.032). For the four individual comparisons the difference in $\boldsymbol{I}_M/\boldsymbol{I}_F$ was significant for one of the two studied populations of *C. maculatus* ($P_{MCMC}$ = 0.006 and 0.32 respectively) and for one of the two populations of *A. obtectus* ($P_{MCMC}$ < 0.001 and 0.14, respectively) (Fig. A3).

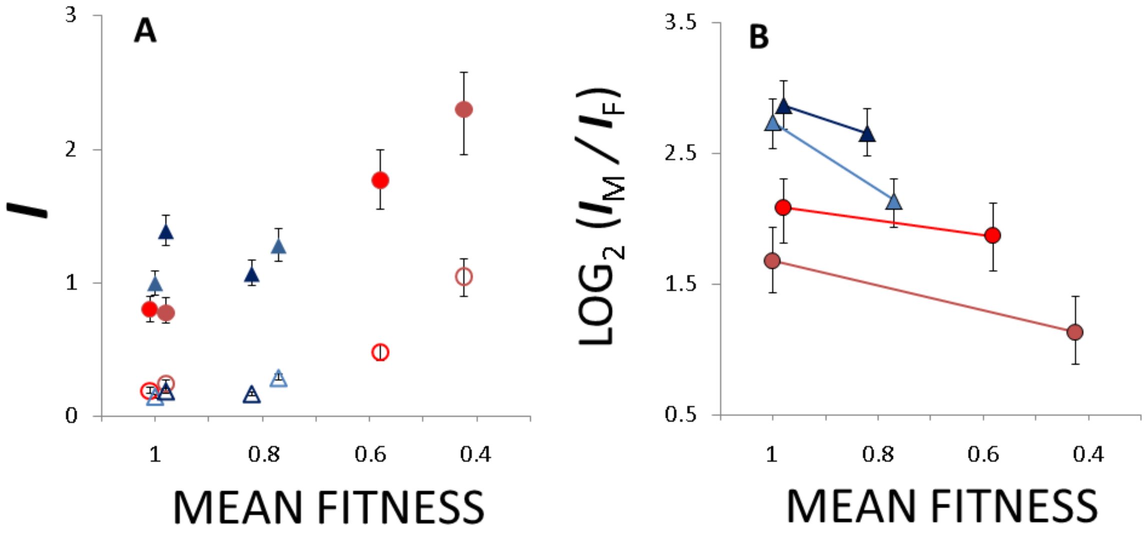


**Fig. A3**. (A) $\boldsymbol{I}_M$ and $\boldsymbol{I}_F$ and (B) their ratio $\boldsymbol{I}_M/\boldsymbol{I}_F$ are plotted against mean fitness averaged across male and female assays as a measure of stress in each assay environment. The fitness in the benign environment is standardized to 1 for each case. In (A) $\boldsymbol{I}_M$ is marked with filled symbols and $\boldsymbol{I}_F$ with open symbols. The two populations of *C. maculatus* are denoted by red colored circles and the two populations of *A. obtectus* are denoted by blue colored triangles.
